# The Noisy Encoding of Disparity Model Predicts Perception of the McGurk Effect in Native Japanese Speakers

**DOI:** 10.1101/2024.04.29.591688

**Authors:** John F. Magnotti, Anastasia Lado, Michael S. Beauchamp

## Abstract

The McGurk effect is an illusion that demonstrates the influence of information from the face of the talker on the perception of auditory speech. The diversity of human languages has prompted many intercultural studies of the effect, including in native Japanese speakers. Studies of large samples of native English speakers have shown that the McGurk effect is characterized by high variability, both in the susceptibility of different individuals to the illusion and in the frequency with which different experimental stimuli induce the illusion. The noisy encoding of disparity (NED) model of the McGurk effect uses Bayesian principles to account for this variability by separately estimating the susceptibility and sensory noise for each individual and the strength of each stimulus. To test whether the NED model could account for McGurk perception in a non-Western culture, we applied it to data collected from 80 native Japanese-speaking participants. Fifteen different McGurk stimuli were presented, along with audiovisual congruent stimuli. The McGurk effect was highly variable across stimuli and participants, with the percentage of illusory fusion responses ranging from 3% to 78% across stimuli and from 0% to 91% across participants. Despite this variability, the NED model accurately predicted perception, predicting fusion rates for individual stimuli with 2.1% error and for individual participants with 2.4% error. Stimuli containing the unvoiced pa/ka pairing evoked more fusion responses than the voiced ba/ga pairing. Model estimates of sensory noise was correlated with participant age, with greater sensory noise in older participants. The NED model of the McGurk effect offers a principled way to account for individual and stimulus differences when examining the McGurk effect within and across cultures.

## Introduction

In the McGurk effect, pairing an auditory syllable with an incongruent visual syllable produces the percept of a third syllable different than either component syllable (McGurk and MacDonald, 1976). The illusion demonstrates the powerful influence of visual information on auditory speech perception and has become a popular instrument for examining audiovisual integration (Beauchamp, 2018).

A number of models have been developed to account for various properties of the McGurk effect. The pioneering fuzzy logic model of Massaro (Massaro, 1998) was developed to explain why some pairings of incongruent audiovisual combinations produce an illusory fusion percept, but for most, participants report the auditory component of the stimulus. More recent models often incorporate principles of Bayesian inference (Andersen and Winther, 2020; Lindborg and Andersen, 2021) and causal inference (Magnotti et al., 2018, 2013; Magnotti and Beauchamp, 2017) although models featuring dynamic predictive mechanisms (Olasagasti et al., 2015) and parallel linear dynamic processes (Altieri and Yang, 2016) have also been proposed.

One challenge to modeling studies of the McGurk effect is the relatively recent realization that there is enormous variability in the McGurk effect across experimental stimuli and participants (Basu Mallick et al., 2015; Jiang and Bernstein, 2011; Stevenson et al., 2012). The original description of the McGurk effect used stimuli recorded from a single talker and reported that the illusion was experienced by nearly all participants. In contrast, Basu Mallick and colleagues tested 12 different McGurk stimuli used in published studies, and found that the efficacy of the different stimuli ranged from 17% to 58%. Across participants, some never perceived the illusion (0%) while others perceived the illusion on every presentation of every stimulus (100%) (Basu Mallick et al., 2015).

The noisy encoding of disparity (NED) model was developed in response to this observation of high variability. Rather than attempt to model the perceptual processes that produce the McGurk effect (as in the models described above), the NED model uses Bayesian, probabilistic inference to predict variation in the McGurk effect. Stimulus differences are modeled using a single parameter for each stimulus (the audiovisual disparity inherent in the stimulus), while participant differences are modelled with two parameters for each participant (an audiovisual disparity threshold, and a sensory noise measure). Using these three parameters, the NED model was able to accurately predict perception in a sample of 165 native English speaking participants (Magnotti and Beauchamp, 2015). NED also accurately modelled perception in 8 adult cochlear implant users and 24 normal-hearing subjects who were native German speakers. The NED model validated the experimental prediction of stronger audiovisual integration in cochlear implant users while controlling for differences between stimuli (Stropahl et al., 2017).

However, both of these studies fit the NED model to participants from Western cultures (native speakers of English and German, respectively). There are many differences between Westerners and East Asians in language and cognition (Varnum et al., 2010). In particular, perception of the McGurk effect has been reported to be markedly reduced in native Japanese speakers (Sekiyama and Tohkura, 1993, 1991). This suggests that applying the NED model to Japanese participants could provide a stringent test of the applicability of the NED model across cultures. To examine this idea, we measured the percepts of 80 native Japanese-speaking participants presented with fifteen different McGurk stimuli.

## Methods

All experiments were approved by the Institutional Review Board of the University of Pennsylvania, Philadelphia, PA.

Data was collected from 101 Japanese-speaking participants recruited by author A.L. from the communities of the Okinawan Institute of Science and Technology, Kyoto University, and the University of Tokyo. All data and analysis code are available in the *Supplementary Material* file. Participants received an Amazon gift card for ¥2000 upon completion of the experiment.

A total of 240 audiovisual stimuli were presented to each participant in pseudorandom order. The stimulus set consisted of 15 McGurk videos, each containing the incongruent pairing of auditory *ba* with visual *ga* (AbaVga, AbabaVgaga) or auditory *pa* and visual *ka* (ApaVka, ApapaVkaka) (McGurk and MacDonald, 1976). The stimulus set was the same as in the original description of the NED model (Magnotti and Beauchamp, 2015) with an additional recording of ApapaVkaka recorded by Arnt Maasø, as described in (Magnotti et al., 2024). There were 8 male talkers and 7 female talkers; two talkers (1 M and 1F) were native Japanese speakers. Full details are provided in the supplementary data file; additional information in *Table 1* of (Basu Mallick et al., 2015) and *Figure 1* of (Magnotti et al., 2015). Each McGurk video was presented 10 times to each participant. As a control, each participant was also presented with 90 congruent audiovisual stimuli (30 different stimuli presented three times each). Each of the 30 congruent stimuli was recorded from a different talker (9 male, 21 female, no overlap with the McGurk talkers). The syllable composition of the congruent stimuli was 5 AbaVba, 5AgaVga, 5 AdaVda, 5 ApaVpa, 5 AkaVka, 5 AtaVta.

**Figure 1.**
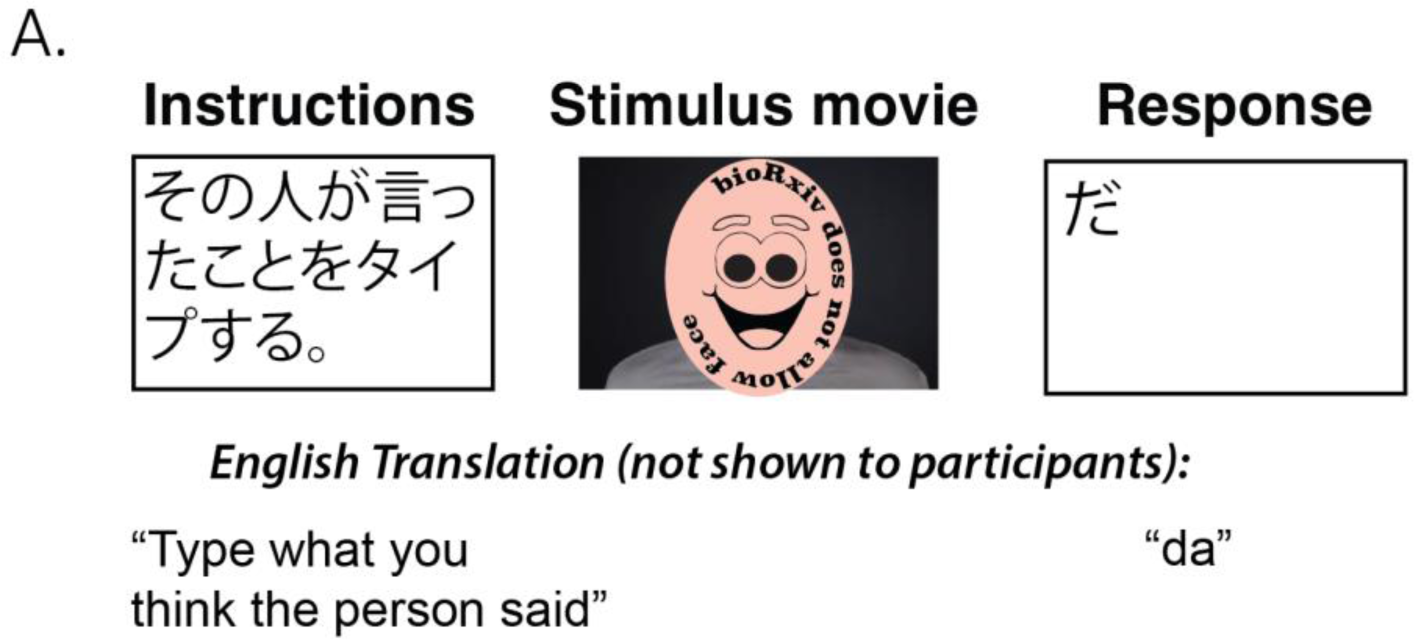
Participants were instructed to report their percept of audiovisual movies containing McGurk and congruent syllables. On each trial, a stimulus movie was played, and participants entered their response using a free-text response box.

### Experimental procedures

The study was conducted online using the SoSci Survey research platform (“Information about SoSci Survey,” n.d.) At the beginning of the experiment, all participants received a short description of the study in Japanese:

様々な音節を発音する人々のビデオが流れます。

この人たちが何を言っているのか、回答してください。

この回答には、約 10 分かかります。

必ず静かな環境で、またはノイズキャンセリングヘッドホンを着用してビデオを見てください。

携帯電話ではなく、必ずノートパソコンまたは PC でビデオを見てください。

Translated as “A video will show people pronouncing different syllables. Please, type what these people are saying. This task will take approximately 10 minutes. Please, watch the videos in a quiet environment or wear noise-cancelling headphones. Please, watch the videos on a laptop or PC, not a mobile phone.“

Participants were also instructed to complete the study on a laptop or a PC. The size of the video was set to a fixed width of 1300 x 650 pixels. Figure 1 shows the structure of each trial. Participants saw the video, an open-choice response box, and the instructions:

ビデオのフレーム全体(上のボックス)が見えるように、ブラウザのウィンドウを調整してください。

上の再生ボタンを押してから、その人が何と言ったか、あなたの考えを下のボックスに入力してください。

間違った答えはありませんので、何度も聴いたり見たりする必要はありません。

Translated as “Adjust your browser window so that you can see the entire frame of the video. Press the play button above and then type what you think the person said in the box below.There are no wrong answers, so please, play the video only once.” The response text box could accept any type of writing (Latin or any traditional Japanese alphabet); the text box could accept any type of writing.

### Data analysis

Fusion responses to the McGurk videos were defined as “da”, “ta”, or “tha”. Responses were translated into English and assigned to one of four mutually exclusive categories: auditory responses, visual responses, fusion responses, and other responses (e.g., “ha”, “va”). For McGurk stimuli consisting of double syllables, each syllable received a half-point rating. For example, the response “dada” was given a score of 1.0 as a complete fusion response, while “bada” was rated as 0.5 for auditory and 0.5 for fusion.

Participants with less than 90% accuracy for congruent syllables identification were excluded from the analysis, ensuring that accuracy for congruent syllables in the remaining participants was high (mean = 96%, standard error of the mean across participants, SEM = 0.3%, range = 90% to 100%). Congruent stimuli almost never evoked fusion responses (2 out of 7200 trials, 0.03%).

Excluding participants with low congruent accuracy left 80 participants whose data are reported in the manuscript. For these participants, 36 self-reported as male and 44 as female with a mean age of 29 (range from 18 to 63). All participants completed high school and 80% had also completed at least a professional or bachelor’s degree. All participants reported their native language as Japanese. Participants were asked to rate their level of English proficiency on a 0 to 6 scale (0 meaning no proficiency at all) with a mean score of 3.0. All participants reported normal hearing and normal or corrected-to-normal vision.

### Noisy encoding of disparity model

The noisy encoding of disparity (NED) model was fit as described in (Magnotti & Beauchamp, 2015). All analysis code is available in the Supplementary file *Full_Results.html*. The model calculates the long-run probability of a fusion percept for each participant and stimulus as

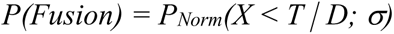

where *X* is the disparity encoded by the participant on an individual trial. The model parameters are *D*, the stimulus disparity; *T*, the participant’s disparity threshold; and *σ*, the participant’s sensory noise level (the standard deviation of the sensory encoding distribution). These parameters were determined by minimizing the absolute error between the model’s predictions and the fusion rate measured for each participant and stimulus. Error was calculated as the mean absolute error (MAE) for individual subjects across stimuli, individual stimuli across subjects, and for each subject-by-stimulus combination.

### Generalization Testing

To assess the generalizability of the model results, a hold-out procedure was implemented. For each participant, the NED model was fit without that subject’s data to obtain stimulus disparity values. Next, holding out a single stimulus, the best-fitting threshold and sensory noise parameters were determined for the held-out subject (without using data from the held-out stimulus). Using the fitted subject parameters and the stimulus disparity values obtained from other participants, the subject’s fusion perception was predicted for the held-out stimulus. This procedure was repeated for each stimulus, resulting in predicted fusion response for each stimulus that was unbiased by the subject’s data for that stimulus.

### Predictors of participant and stimulus variability

Multiple linear regression models were constructed to examine the relationship between participant-level model parameters (one model for disparity threshold, one for sensory noise) and participant demographic variables (age, gender, English proficiency, and highest education level). The models were obtained using stepwise regression to automatically select the best parameters using the Bayesian Information Criterion (BIC) cost function. Because of the small sample size, no interactions were allowed during the stepwise model building procedure. The initial model was the model including all variables, and the minimal model was the intercept-only model.

To understand stimulus-level variation, the same procedure was applied to the stimulus parameter of disparity, with stimulus variables of syllabic content (voiced AbaVga *vs*. voiceless ApaVka), talker gender (male *vs*. female) and talker native language (Japanese *vs*. non-Japanese).

## Results

### Responses to McGurk stimuli

Across stimuli, participants reported an average of 23% fusion responses. There was a high degree of variability in the number of fusion responses across the 15 different McGurk stimuli: the weakest stimulus evoked the McGurk effect on 3% of trials while fusion responses were evoked on 78% of trials by the strongest stimulus. There was also a high degree of variability in the percentage of McGurk responses across different participants: across stimuli, the least-susceptible participant perceived the illusion on 0% of trials while the most-susceptible participant perceived the illusion on 91% of trials. The combination of high stimulus and high participant variability meant that for 11 of the 15 McGurk stimuli, the frequency of fusion responses ranged from the lowest possible value of 0% for some participants to the highest possible value of 100% for other participants.

### Model fitting examples

Figure 2A illustrates the model fitting process for three stimuli which evoked different average rates of fusion perception. For each stimulus, the model estimates an audiovisual disparity, which is assumed to be constant (*i.e.* a physical property of the stimulus). On each presentation of a stimulus, a participant is assumed to sample the audiovisual disparity of the stimulus. Due to sensory noise, the measurement is not precise, resulting in a Gaussian distribution of disparity estimates across presentations. The mean of the distribution is centered on the true disparity of the stimulus, and the degree of sensory noise (width of the Gaussian distribution) is estimated by the model separately for each participant. For each stimulus presentation, the participant compares the estimated the stimulus disparity with an internal threshold. If the estimated stimulus disparity exceeds their internal threshold, the participant assumes that the auditory and visual component of the speech come from different talkers, and perception defaults to the auditory component of the stimulus. If the estimated disparity does not exceed their threshold, the participant integrates the auditory and visual components of the stimulus and perceives the illusion.

**Figure 2.**
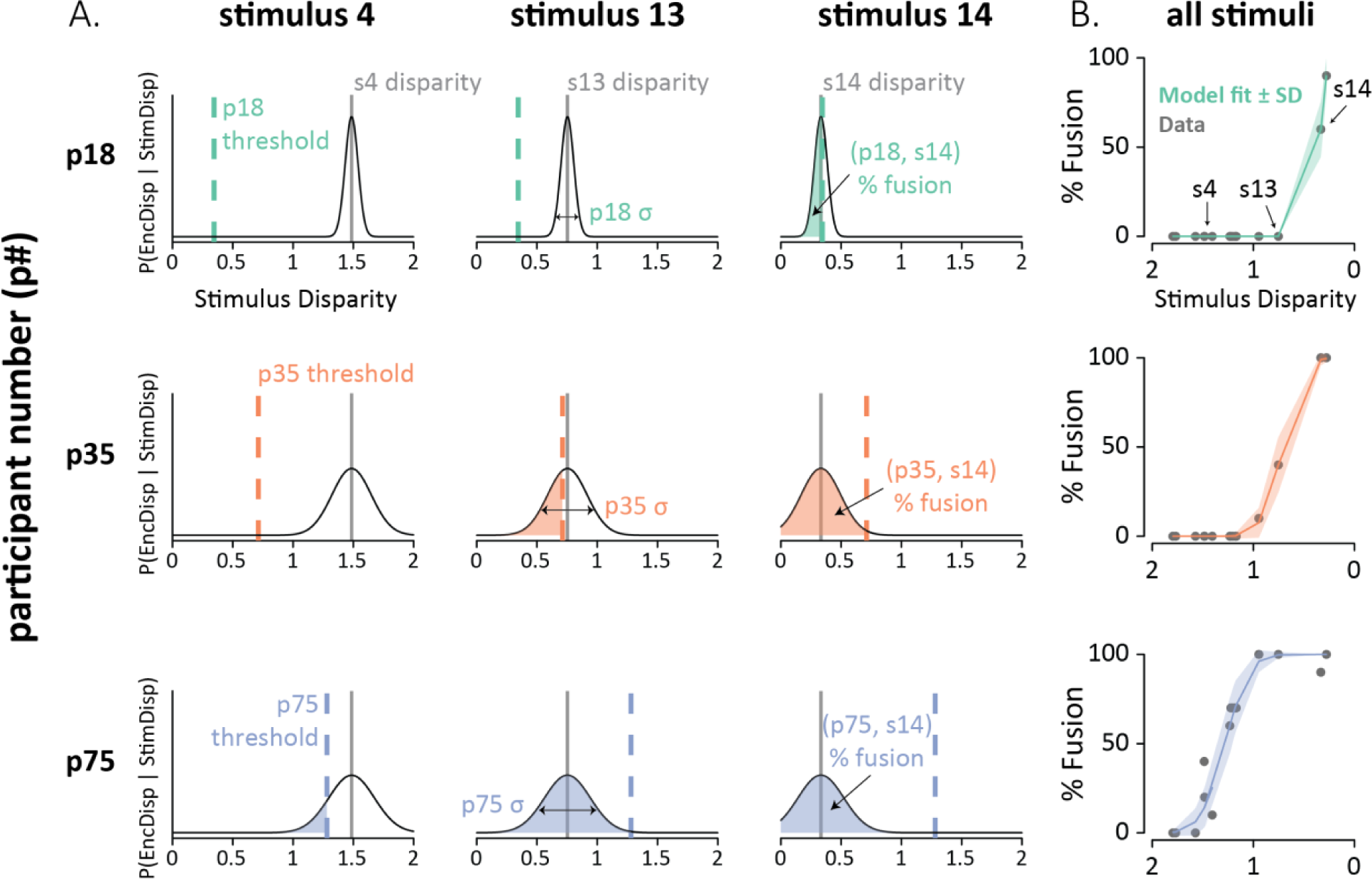
**A.** Fits of the noisy encoding of disparity (NED) model across participants and stimuli. Each row shows a single participant (p18, p35, p75; one color per participant). Each column shows a different stimulus (s4, s13, s14). The x-axis shows the estimated stimulus disparity. The y-axis shows probability. The thin black line is the Gaussian probability distribution of the disparity estimates. The stimulus disparity is fixed for each stimulus (gray vertical lines). On each presentation of a stimulus, the stimulus disparity is estimated by the observer with sensory noise, σ (horizontal arrows; fixed for each participant). The estimated disparity is compared to the participant’s integration threshold (vertical colored dashed line, fixed for each participant). If the estimated disparity is below the threshold, then the participant experiences the McGurk fusion percept. The shaded area is the predicted percent fusion for that participant and stimulus. **B.** Summary of the model fit for all stimuli for the three participants. The x-axis shows the estimated stimulus disparity, with values reversed to produce an increasing psychometric function. The y-axis shows the % fusion reports for each stimulus. Each gray point shows data, colored line shows model prediction (shaded color region shows model SD).

For participants with three very different rates of average fusion perception: low (p18; 10%), moderate (p35; 17%) and high (p75; 53%), the model accurately predicted perception across the 15 different McGurk stimuli (Figure 2B). Note that simply predicting the average fusion rate for each participant would produce much larger errors than the predictions made by the model. For instance, for p18, stimulus 14 evoked 60% fusion percepts, exactly the same as the model prediction (0% error) while the mean fusion rate for p18 was 10% (50% error if used as a prediction). For p75, stimulus 1 produces 0% fusion percepts, very close to the model prediction of 0.3% (0.3% error). In contrast, the mean fusion rate for p75 was 53% (53% error if used for prediction). Similarly large errors result if the mean fusion rate for each stimulus is used for prediction.

### Model fitting

Figure 3 shows the model results across stimuli and participants. The model accurately reproduced stimulus-level variation, predicting stimulus-level mean fusion responses with a mean absolute error of 2.1% ± 0.4% SEM). The model reproduced subject-level variation with an average error of 2.4% ± 0.3%. For single stimulus-participant pairs, the response could be predicted with an average error of 4.7% ± 0.5%.

**Figure 3.**
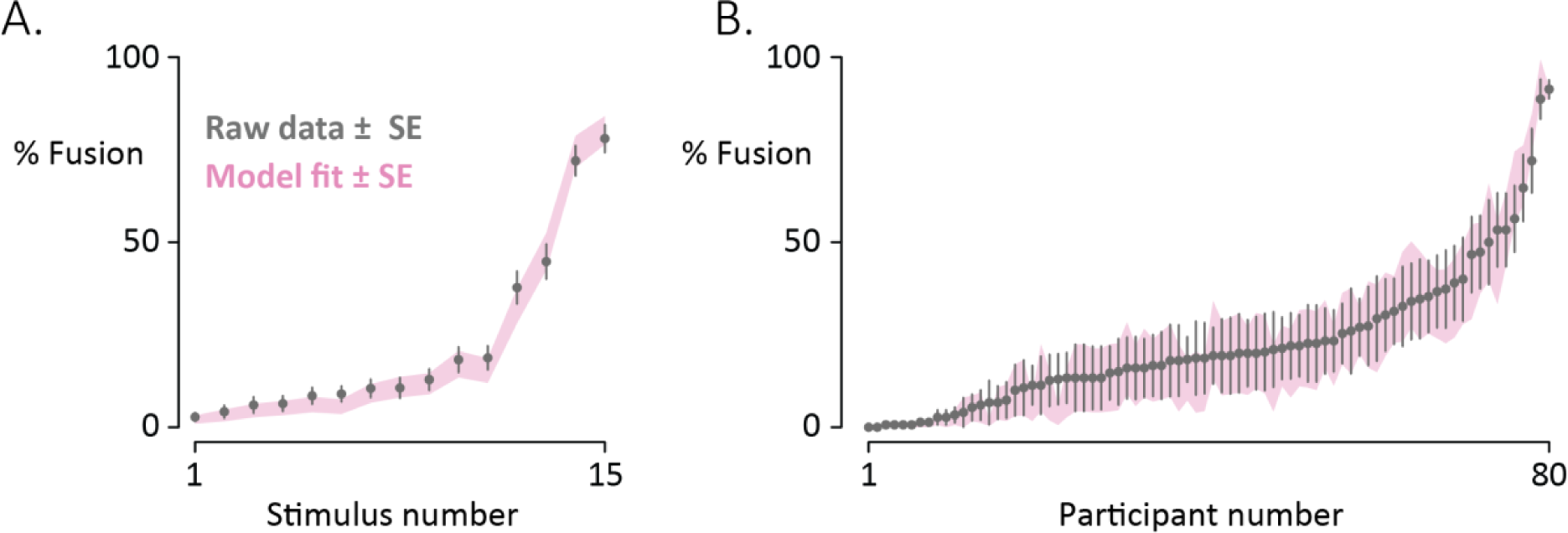
**A.** Participants reported their percepts of 15 different McGurk stimuli (10 repetitions each, randomly interleaved with congruent speech). Gray circles denote the mean percentage of fusion responses across participants for each McGurk stimulus (bars show standard error of the mean across participants.) Stimuli are sorted from fewest to most fusion responses. The pink shaded region shows the fit of the noisy encoding of disparity model (mean ± one standard error). **B.** For each of 80 participants, the mean percentage of fusion responses across the 15 different McGurk stimuli was calculated (one gray circle per participant; bars show standard error of the mean across stimuli; participants sorted from fewest to most fusion responses). The shaded region shows the fit of the noisy encoding of disparity model (mean ± one standard error).

Validating the model assumption of a constant audiovisual disparity for each stimulus, the stimulus ranks were highly correlated across subjects (average subject-level rank correlation with global rank, *r* = 0.68 ± 0.02, *p* < 10^−16^).

We also estimated out-of-sample generalization using leave-one-stimulus-out model fitting. Fusion percentages on untrained stimuli could be predicted with an error of 9.7% ± 0.5%. The error rate was low for untrained stimuli (9.7% *vs*. 4.7% for trained data) demonstrating that the high accuracy of the model predictions was not due to overfitting the training data.

### Stimulus Differences

The 15 McGurk stimuli differed along several dimensions, including syllabic composition (the voiced syllables AbaVga *vs.* the voiceless syllables ApaVka); the gender of the talker; and the native language of the talkers (Japanese *vs.* non-Japanese). To test the importance of these factors to the NED model, we used stepwise linear regression to find the best model of audiovisual stimulus disparity as a function of syllable, talker gender, and talker native language. The best-fitting model explained 70% of the variance in the parameter [*R*^2^ = 0.70, *F*(1, 13) = 30.2, *p* = 10^−4^] and included only syllable content (*b* = −0.92). The mean fusion rate was 65% for voiceless ApaVka stimuli compared with 12% for voiced AbaVga stimuli.

A possible concern is confounding of talker differences and syllable content. This concern was mitigated by the fact that the stimulus set contained examples of both pairings recorded from the same talkers (Magnotti et al., 2024). For talker Audrey Nath, the pa/ka pairing evoked 72% fusion responses while ba/ga evoked 6% fusion responses. For talker Arnt Maasø, pa/ka evoked 45% fusion while ba/ga evoked 11% fusion responses.

Neither talker gender [*F*(1, 11) = 1.2, *p* = 0.3] nor talker language [*F*(1, 11) = 10^−4^, *p* = 0.99] were significant predictors of stimulus-level differences.

### Participant Differences

The NED model estimated sensory noise and disparity threshold parameters for each participant. Statistical modeling revealed two significant associations between these parameters and participant demographic variables. There was a significant positive correlation between the sensory noise parameter and age of the participant [*b* = 0.003, *R*^2^ = 0.067, F(1,78) = 5.6, *p* = 0.02] indicating greater sensory noise with increasing age. In contrast, the disparity threshold parameter was not predicted by participant age [*b* = 0.002, R^2^ = 0.003, F(1,78) = 0.20, *p* = 0.66].

Conversely, there was a significant negative correlation between self-reported English proficiency rating and the disparity threshold parameter [*b* = −0.07, R^2^ = 0.066, F(1,78) = 5.5, *p* = 0.02] while English proficiency was not predictive of sensory noise [*b* = −0.012, R^2^ = 0.026, F(1,78) = 2.1, *p* = 0.15].

## Discussion

The noisy encoding of disparity (NED) model was fit to the perceptual reports of native Japanese speakers presented with McGurk stimuli. Despite the high variability of the McGurk effect (ranging from 0% to 100% fusion reports across participants for most stimuli), the NED model predicted perception of the illusion with only a few percent error for individual stimuli, participants, and stimulus-participant pairs.

The NED model makes two fundamental assumptions. First, that individual differences in audiovisual speech perception can be captured by two simple parameters, that of sensory noise and sensitivity to audiovisual disparity. Second, that different McGurk stimuli can be characterized by the amount of audiovisual disparity they contain. The ability of the NED model to accurately predict perception in native Japanese speakers, in native English speakers, native German speakers, and native German adults with cochlear implants (Magnotti and Beauchamp, 2015; Stropahl et al., 2017) suggest that these assumptions are justified.

To more concretely test the assumption that different McGurk stimuli can be characterized by an intrinsic audiovisual disparity, we compared the model results for the present study and original description of the model, which used identical stimuli (with the exception of one additional stimulus in the current study; data from this stimulus were not considered). Comparing stimulus ranks between native Japanese and native English speakers revealed a high correlation, *r* = 0.87, *p* < 10^−16^ demonstrating the reasonableness of defining an intrinsic disparity for each stimulus.

### Participant variability and intercultural comparisons

In the present study, we found high variability in the McGurk effect across native Japanese speakers, consistent with the high variability observed in studies of native English speakers (Basu Mallick et al., 2015; Jiang and Bernstein, 2011; Stevenson et al., 2012). The high variability inherent in the McGurk effect *within* single cultures complicates studies of potential differences in the effect *across* cultures. Using simulations, Magnotti and Beauchamp (Magnotti and Beauchamp, 2018) estimated the number of participants required to detect group differences in the McGurk effect with 80% power, a common statistical benchmark (Cohen, 1992). Even assuming a moderate mean difference of 10% in fusion rate across groups, more than 300 participants would be required to reliability detect the difference.

The large sample size required for well-powered detection of group differences in fusion rates necessitates careful evaluation of published studies: a “statistically significant” finding in an underpowered study may greatly inflate the measured effect-size (Gelman and Weakliem, 2009). Intercultural comparisons of the McGurk effect with larger sample sizes have largely failed to detect any difference in fusion rates. A study with a sample size of 307 did not find a significant difference in fusion rates between native English speakers tested in the U.S.A. and native Mandarin speakers tested in China (Magnotti et al., 2015). A study with a sample size of 99 did not find a significant difference in fusion rates between native Finnish speakers and native Japanese speakers (Tiippana et al., 2023). In contrast, a study reporting low rates of McGurk fusion in native Japanese speakers tested just 10 participants (Sekiyama and Tohkura, 1991).

### Stimulus variability

Just as variability across participants complicates intercultural comparisons, so does variability across stimuli. Across the fifteen different McGurk stimuli tested in the present study, there was high variability in the rate of fusion percepts, ranging from 3% to 78%. This large variability is problematic when making inferences from only a few stimuli.

For instance, there are mixed reports in the literature about the existence of cross-language influences in the McGurk effect. Some studies report *more* fusion responses when native speakers of one language are presented with McGurk stimuli recorded by a native talker of another language (Ujiie and Takahashi, 2022); other studies report *fewer* fusion responses or mixed results (Chen and Hazan, 2007). In the present study, there was no significant difference in fusion responses between the two stimuli recorded by native Japanese speakers and the remaining stimuli, recorded by native speakers of other languages.

However, given the high variability across stimuli recorded by talkers of the same native language, or even multiple stimuli recorded from the same talker (Magnotti et al., 2020), attempts to identify differences between stimulus categories with only one or two examples from each category are unlikely to yield reliable results. Instead, it is important to test as many stimuli from each category as possible. The largest study in this vein tested Japanese and Finnish participants using McGurk stimuli recorded by four native Japanese talkers and four native Finnish talkers and found no significant difference between Japanese and Finnish talkers (Tiippana et al., 2023).

### Limitations

A limitation of the NED model is that it is primarily descriptive rather than mechanistic. For instance, it takes as a given that participants who integrate of auditory “ba” with visual “ga” report the fusion percept of “da”, rather than providing an explanation for the fusion percept. More mechanistic models often ignore individual differences but incorporate sequential steps of unisensory estimation and multisensory integration using principles of Bayesian inference (Andersen and Winther, 2020; Lindborg and Andersen, 2021) and causal inference (Magnotti et al., 2018, 2013; Magnotti and Beauchamp, 2017). Fitting these more mechanistic models requires substantially more data, including perceptual measurements of unisensory auditory and visual speech, usually with sensory noise added to each modality at varying levels or different degrees of temporal asynchrony between modalities. In contrast, the simpler NED model only requires measuring the perception of McGurk stimuli.

It remains to be clarified how the parameters in the NED model relate to real-world variables. The model assumes that audiovisual disparity is an intrinsic property of different McGurk stimuli. The use of advanced synthetic faces should also allow more insight into understanding and manipulating the factors contributing to stimulus disparity (Shan et al., 2022; Thézé et al., 2020; Varano et al., 2022; Yu et al., 2024), as should measurements of the mouth and face movements made by real talkers (Jiang et al., 2007). The NED model fits a sensory noise parameter for each participant, with the finding that sensory noise increases with age. The variability of the BOLD fMRI response to audiovisual speech also increases with age (Baum and Beauchamp, 2014). This suggests that measuring neural variability in speech processing regions could allow an independent assessment of sensory noise, linking a NED model parameter to brain activity. A recent fMRI study found that observers’ response entropy was greater for McGurk compared to congruent audiovisual stimuli, corresponding to increased BOLD activity in brain regions important for cognitive control (Dong et al., 2024). Parietal and frontal regions are important for causal inference on audiovisual stimuli (Gau and Noppeney, 2016; Mihalik and Noppeney, 2020). Brain activity in these regions could be measured to provide an independent estimate of a participant’s disparity threshold for integrating auditory and visual speech.

## Notes

### Competing Interest Statement

The authors have declared no competing interest.

